# Linkage mapping and QTL analysis of flowering time using ddRAD sequencing with genotype error correction in *Brassica napus*

**DOI:** 10.1101/2020.06.26.162966

**Authors:** Armin Scheben, Anita Severn-Ellis, Dhwani Patel, Aneeta Pradhan, Stephen Rae, Jacqueline Batley, David Edwards

## Abstract

*Brassica napus* is an important oilseed crop cultivated worldwide. During domestication and breeding of *B. napus*, flowering time has been a target of selection because of its substantial impact on yield. Here we use double digest restriction-site associated DNA sequencing (ddRAD) to investigate the genetic basis of flowering in *B. napus*. An F_2_ mapping population was derived from a cross between an early-flowering spring type and a late-flowering winter type. Flowering time in the mapping population differed by up to 25 days between individuals. High genotype error rates persisted after initial quality controls, as suggested by a genotype discordance of ∼12% between biological sequencing replicates. After genotype error correction, a linkage map spanning 3,605.70 cM and compromising 14,630 single nucleotide polymorphisms (SNPs) was constructed. A quantitative trail locus (QTL) on chromosome C2 was detected in the vicinity of flowering time genes including *FT* and *FLC*. These findings demonstrate the effectiveness of the ddRAD approach to sample the *B. napus* genome. Our results also suggest that ddRAD genotype error rates can be higher than expected in F_2_ populations. Quality filtering and genotype correction and imputation can substantially reduce these error rates and allow effective linkage mapping and QTL analysis.

## Introduction

Genotyping-by-sequencing (GBS) is a powerful tool for high-throughput discovery of genetic polymorphisms in crops (*1-5*). GBS comprises a range of library preparation and sequencing approaches that differ in their costs, methodical biases and the type and amount of data produced (*1, 6*). Restriction site-associated DNA sequencing (RAD) is a GBS method that can be used to cost-effectively calibrate the number and coverage of genotyped loci and single nucleotide polymorphisms (SNPs) by varying the enzymes used and the sequencing depth. A recent comparative analysis of single enzyme RAD and two enzyme double digest RAD (ddRAD) used a range of enzyme combinations in different plants and suggested that the enzyme combination of HinfI and HpyCH4IV was promising for maximising genome coverage breadth across a range of species (*7*).

GBS has been used for marker discovery, linkage mapping and QTL analysis in a range of crops (*8-10*), including the important oilseed crop *Brassica napus* (*11-13*). Over 20 high density linkage maps have been generated for *B. napus* using RNA sequencing (*14*), the Brassica 60K genotyping array (*15, 16*) and ddRAD sequencing (*12*). Combined with phenotypic data, these linkage maps provide a powerful basis for identification of genes underlying agronomic traits, which can then be introduced into crop germplasm (*17, 18*). Crop yield in *B. napus* is strongly dependent on flowering time, making this trait a key breeding target.

Flowering time genetic pathways have been elucidated in Arabidopsis and most flowering time genes are known to be conserved between Arabidopsis and *B. napus* (*19-21*). Many QTL and associated SNPs for flowering time have been detected in *B. napus* (*22-29*). Despite this progress in understanding the genetic underpinnings of *B. napus* flowering time, a substantial proportion of flowering time variation remains to be explained.

There are three *B. napus* oilseed rape (OSR) growth types with considerable variation in flowering time: spring, semi-winter and winter. Spring OSR and semi-winter OSR have a low requirement for vernalization to flower and are early-flowering, whilst winter OSR has a strong vernalization requirement and is late-flowering. In *B. napus* breeding, the flowering traits of spring OSR decrease its generation time compared to winter OSR, allowing more rapid breeding cycles. Reducing vernalization requirements in winter OSR by introducing spring OSR alleles would be one approach to allow breeders to accelerate winter OSR breeding. In addition, *B. napus* hybrids are generally higher yielding than open pollinated varieties due to heterosis (*30, 31*). If flowering time can be efficiently managed, heterosis could be exploited from hybridization of spring OSR and winter OSR. Identifying flowering time loci that distinguish spring OSR and winter OSR therefore has important breeding applications. Here, to identify these loci, we crossed a spring OSR and a winter OSR to generate an F_2_ mapping population. We genotyped the progeny and parental lines using ddRAD sequencing. Finally, we constructed a high-density linkage map and carried out QTL analysis of flowering time and the related trait budding time. We present candidate regions for flowering time and budding time and discuss the use of error-prone ddRAD genotyping in heterozygous breeding populations.

## Results

### Pre-processing and aligning sequencing reads

A single sample (“146”) was excluded from further analysis as it had fewer than one million reads after trimming. In the remaining 206 samples, a mean of 13.14 million raw paired sequences were generated per sample. A mean of 56.2% of reads were uniquely aligned with high quality. The mean coverage depth at covered bases was 9.41 × and the mean coverage breadth of the genome was 18.03%.

### SNP filtering and genotype correction

A total of 4,841,931 biallelic SNPs were identified in the mapping population and parental individuals. For further analysis, the seven parental individuals were excluded from the SNP set. Filtering by individual missingness, genotype depth, minor allele frequency (MAF), and genotype missingness reduced the number of SNPs to 124,804. Of the 199 progeny, 192 were retained after filtering individuals with high genotype missingness. Of the 124,804 SNPs, 50,856 did not have a heterozygous genotype in any parental individual. The SNPs with heterozygous genotypes in the parental individuals may be caused by mismapping or remaining heterozygosity in the parental genomes and were therefore excluded. Next, removing 16,647 SNPs that were monomorphic between parents and 5,957 that showed segregation distortion (p < 0.01), generated a set of 28,252 SNPs. Segregation distorted SNPs were distributed relatively evenly across chromosomes, with noticeable hotspots at the ends of chromosomes A1 and C5 (Figure S1). Genotype-Corrector quality control removed 13,509 further SNPs after filtering homozygous SNPs located within heterozygous regions. A total of 4.94% of genotypes were corrected using Genotype-Corrector and 94.76% of missing genotypes were imputed (Figure 1). The most frequent genotype corrections were B to AB (29.56%) and A to AB (23.48%).

**Figure 1.**
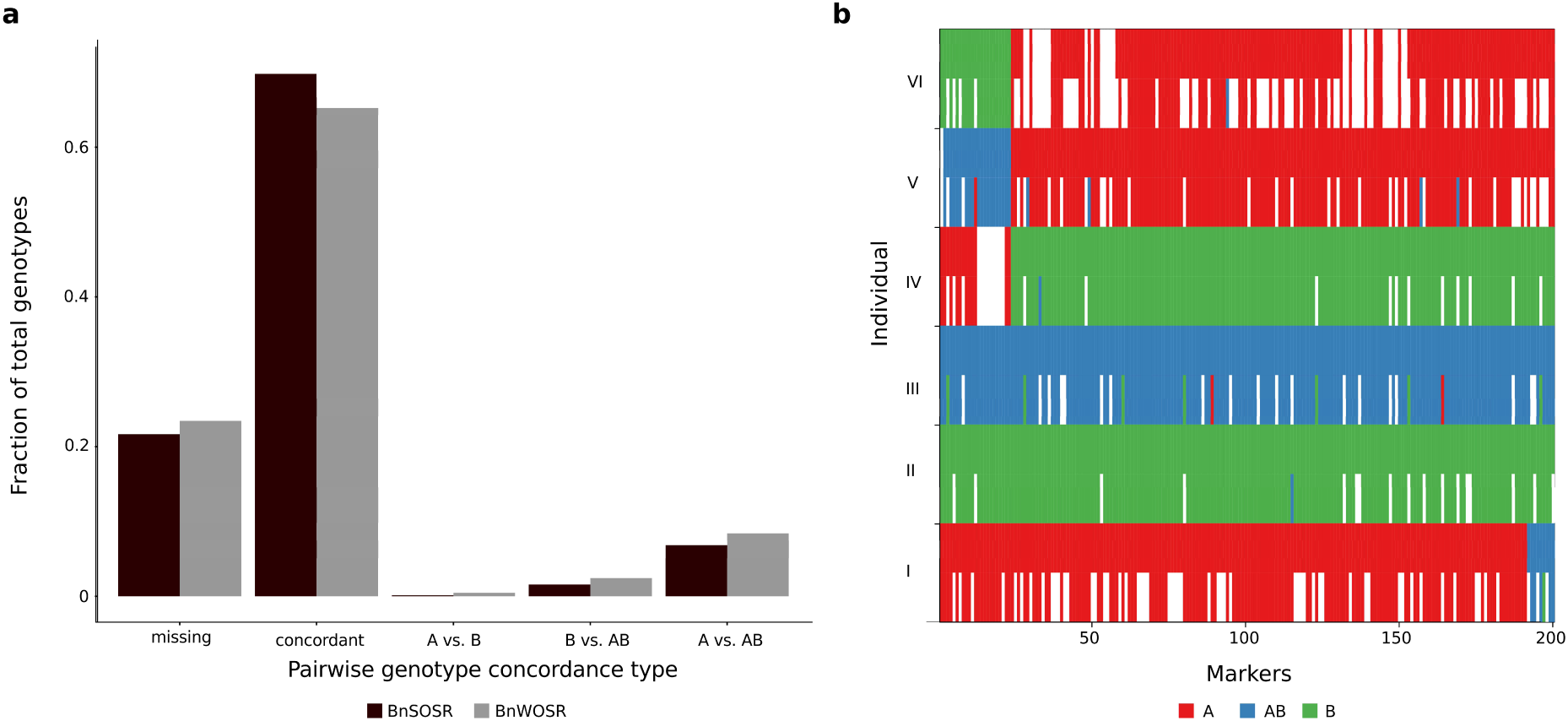
Genotyping errors and correction process. a) Mean pairwise genotype concordance between replicate individuals for the parental lines. The bars for the category ‘missing’ include all genotype pairs where at least one pair had a missing genotype call. Genotypes shown are BnSOSR as ‘A’, BnWOSR as ‘B’ and heterozygous as ‘AB’. b) Comparison of corrected and uncorrected genotypes. An example of genotype correction and imputation using 200 SNPs on chromosome A3 is given for six representative individuals denoted as I-VI (samples 1, 100, 102, 103, 104, and 105). Genotypes are encoded in three colours (A: red; B: green; AB: blue) and missing markers are shown in white. For each individual, the lower row shows the genotypes before the correction and imputation step and the upper row shows the genotypes after this step.

In the parental replicate individuals, analysis of pairwise genotype concordance identified a mean genotype discordance of 12.28% (Figure 1). Discordance between homozygous genotypes (A vs. B) was rare, with conflicts between homozygous and heterozygous genotypes (A vs. AB, B vs. AB) making up 97.51% of genotype discordance.

### Linkage mapping

A linkage map spanning 3,605.70 cM and compromising 14,630 markers was constructed using ASMap with the corrected and imputed markers (Figure 2 and Table S2). The A genome map was 1,771.53 cM with 8,587 markers and the C genome was 1,661.98 cM with 6,043 markers. The highest mean marker density was found on chromosome A10, with 47.90 markers per Mb. Mean marker density per Mb was higher on the A genome (28.90) than the C genome (12.18).

**Figure 2.**
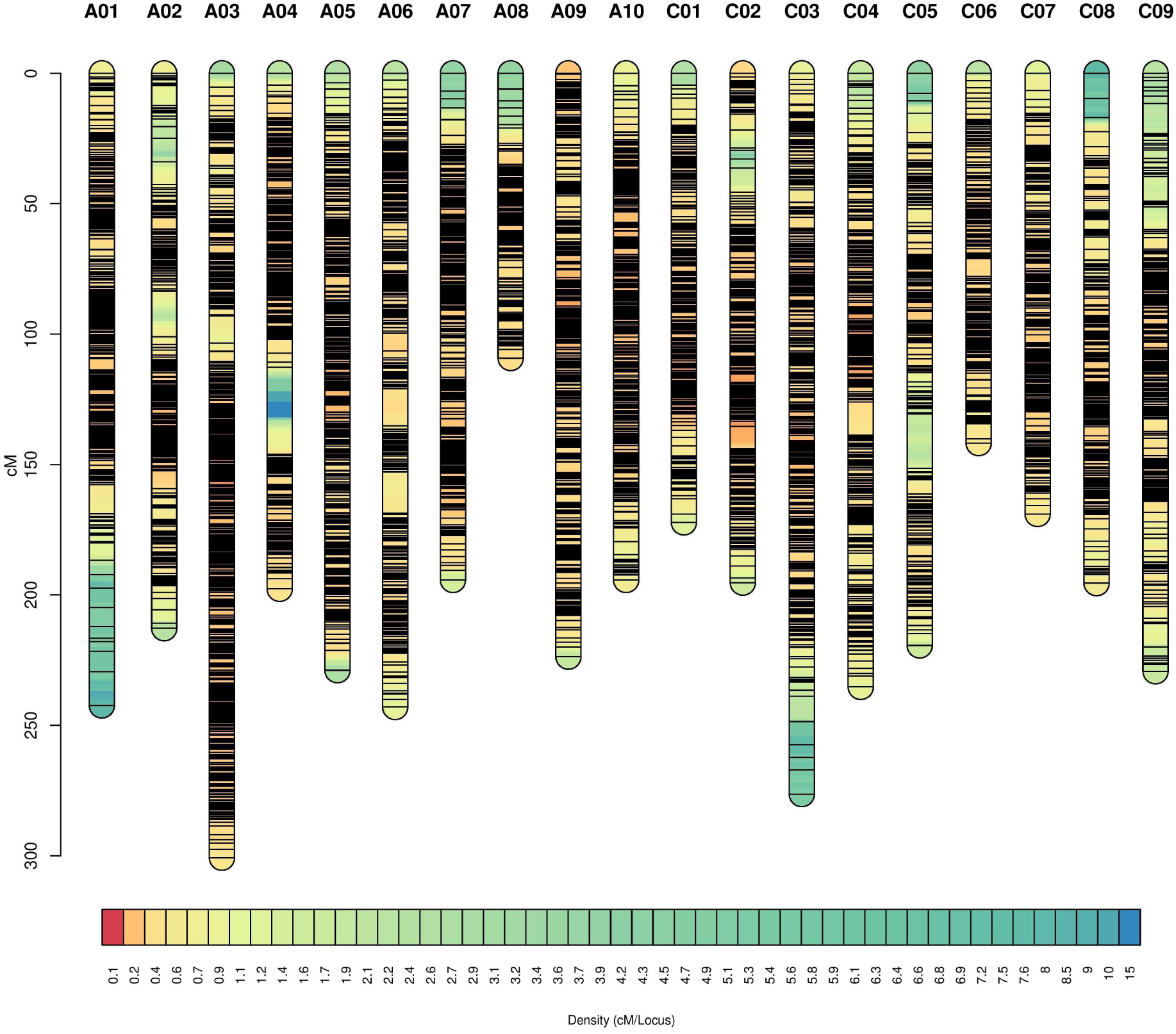
Genetic map based on ddRAD markers. Black lines indicate marker positions and colours along the chromosomes indicate markers density, with extremely dense regions appearing as black due to tightly clustered markers. Figure rendered using LinkageMapView 2.1.2 (*32*).

A supplementary map was constructed using uncorrected markers, which showed a high inflation of genetic distances with a total map length over 30,000 cM (Figure S2). Compared to six published genetic maps that were generated with different approaches, the corrected genetic map still showed some indications of inflation (Table S3).

The correlation between genetic and physical map order provides information about the consistency between the genetic map and the reference genome. The mean Spearman’s rank correlation for marker order per chromosome was 1.0 (Table 1). Several minor inconsistencies in marker order were observed (Figure S3). All chromosomes showed mean correlations over 0.98. Mean individual crossover frequency per chromosome was 2.79.

**Table 1.** Summary of the genetic map. Spearman’s rank correlation (rho) was calculated for the genetic marker positions and the physical marker positions on the reference genomes.

| Chr | Length (Mb) | Length (cM) | Number of markers | $\rho$ |
| --- | --- | --- | --- | --- |
| A1 | 31.16 | 242.40 | 740 | 0.98 |
| A2 | 31.34 | 185.10 | 630 | 1.00 |
| A3 | 39.49 | 300.82 | 1,581 | 1.00 |
| A4 | 23.31 | 58.35 | 748 | 1.00 |
| A5 | 28.6 | 228.92 | 892 | 1.00 |
| A6 | 31.9 | 34.53 | 729 | 1.00 |
| A7 | 28.9 | 194.30 | 954 | 1.00 |
| A8 | 21.74 | 109.17 | 365 | 1.00 |
| A9 | 46.72 | 223.68 | 990 | 1.00 |
| A1 | 19.96 | 194.26 | 958 | 0.99 |
| C1 | 47.95 | 172.18 | 739 | 1.00 |
| C2 | 58.66 | 195.33 | 832 | 0.99 |
| C3 | 71.85 | 276.37 | 900 | 1.00 |
| C4 | 61.04 | 235.26 | 861 | 0.99 |
| C5 | 52.72 | 219.41 | 483 | 1.00 |
| C6 | 44.61 | 141.83 | 504 | 1.00 |
| C7 | 52.5 | 168.96 | 612 | 1.00 |
| C8 | 46.29 | 195.51 | 569 | 1.00 |
| C9 | 60.21 | 229.31 | 543 | 1.00 |
| Total | 798.95 | 3,605.70 | 14,630 | - |

### QTL mapping

BnSOSR flowered 20 days earlier (range: 10 to 28) and went to bud 17 days earlier (range: 12 to 20) on average than winter type BnWOSR (Table S4). In the F_2_ progeny, flowering times were distributed within the parental range (Figure 3). A single significant (p<0.05) overlapping QTL region for the traits flowering time and budding time was detected on chromosome C2 (Figure 3). The physical region of the QTL spanned 20.57 Mb for flowering time and 0.77 Mb for time to bud (Table S5). The flowering time QTL contained 10 flowering time homologs including *FT* (Table S6), and the budding time QTL contained no known flowering time homologs. Carrying the BnWOSR allele at the QTL peak SNP led to an increase in the days to bud and flower (Figure S4). The percentage of phenotypic variance explained for the identified QTL was 9.08% for flowering time and 8.08% for budding time. Suggestive LOD peaks are also noticeable on A2, A3 and C9

**Figure 3.**
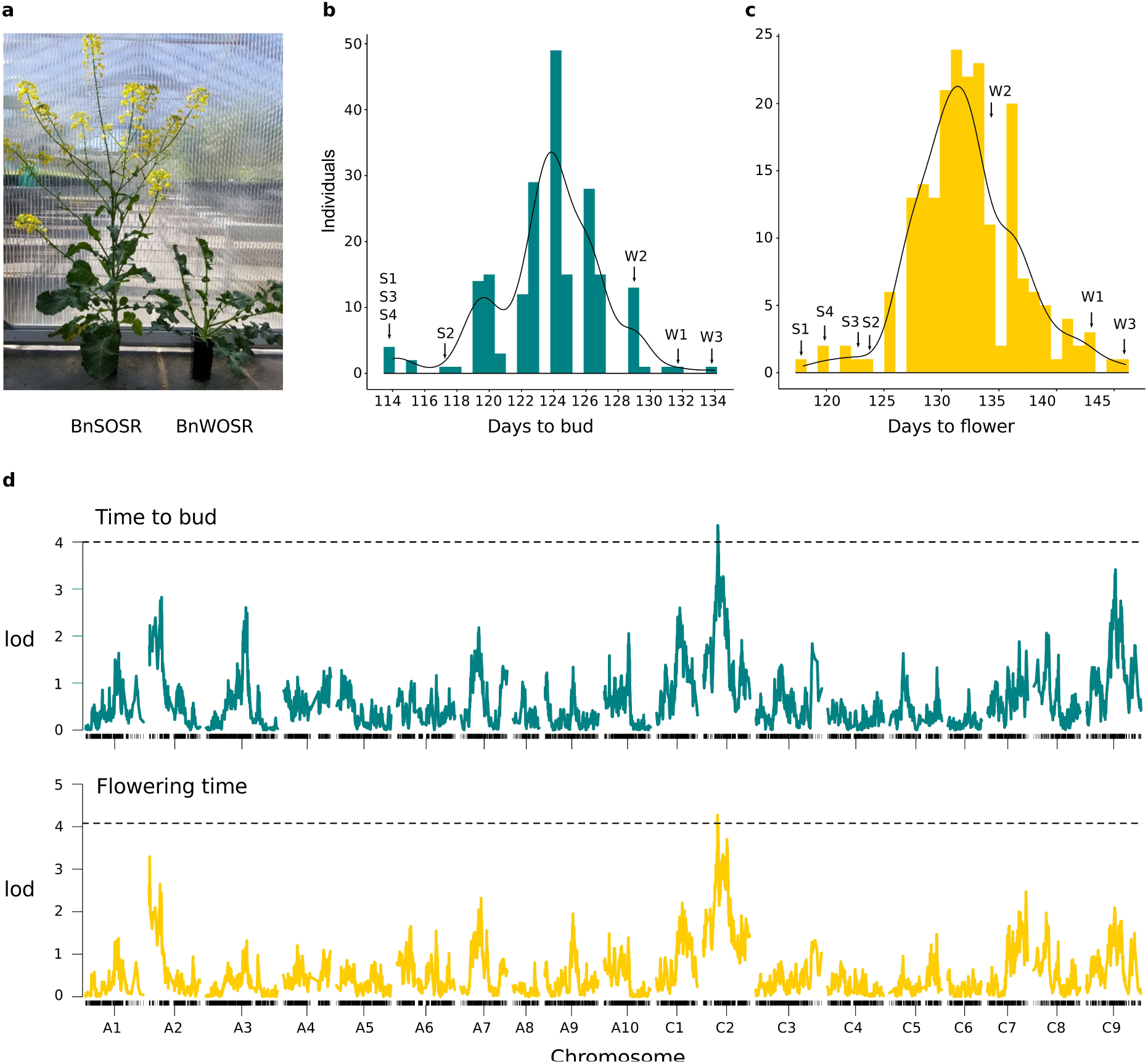
Phenotyping and QTL mapping results. a) A photograph comparing development in parental individuals 126 six days after sowing. b) Histogram of mapping population phenotypes for budding time. c) Histogram of mapping population phenotypes for flowering time. Parental individuals S1-S4 of BnSOSR and W1-W3 of BnWOSR are indicated. Phenotype data is provided in Table S7. d) Genome-wide LOD plots for single QTL mapping of flowering time and budding time. Significant (p<0.05) thresholds based on permutation testing shown with broken lines.

## Discussion

### Optimising genotyping-by-sequencing strategies

Genotyping with ddRAD was effective at generating a set of 14,630 high-quality SNPs for linkage and QTL mapping. The findings of this study can help calibrate the number of sequencing reads and genomic loci required for a range of research goals in *B. napus* and related species. For GBS, researchers often aim to optimize the genome coverage by controlling the ratio of reads sequenced to the number of loci generated. Here, the expected maximum genome coverage breadth based on in silico digestion with the enzymes HpyCH4IV and HinfI was 24.6%. However, the observed mean genome coverage breadth was lower at 18.03%. An even greater inconsistency was reported when using HpyCH4IV and HinfI in Arabidopsis and *Glycine max*, where the expected genome coverage breadth was 29.4% and 23.1% but the observed experimental values were 4.45% and 3.33% respectively (*7*). The inconsistency has been explained as a product of fragment size selection bias, redundant in silico loci, and insufficient sequencing reads (*7*). In the *B. napus* population used here, the most important factor determining coverage was the amount of sequencing reads available for each sample. Indeed, 24 samples with high sequencing effort showed genome coverage breadth greater than the expected 24.6% up to a maximum value of 36.68%. This is particularly surprising, as coverage was calculated based only on reads aligned with high quality, which is expected to substantially reduce coverage breadth. These findings suggest that, at least in *B. napus*, igCoverage can underestimate the maximum achievable genome coverage breadth.

The high genome coverage breadth achieved using HpyCH4IV and HinfI indicates that these enzymes are well suited for high-density sampling of genome-wide diversity in *B. napus*. However, when sequencing effort is uneven between samples, high genome coverage breadth can increase genotype missingness through allele dropout. If a locus is not sequenced in enough individual samples (here the cut off was 50%) at sufficient depth, it is removed during SNP calling or SNP filtering and becomes a missing genotype call. High levels of missingness are a common characteristic of reduced representation sequencing (*3*) and can limit the usefulness of genotype data in studies where genotype imputation is not possible (*33*). Nevertheless, as shown in this study, by combining imputation with a high-density sampling of the genome, the limitations of genotype missingness in a mapping population can be overcome.

### Linkage mapping

The correlation of physical and genetic maps was high, indicating that the map is accurate and collinear with the reference genome. Similarly, collinear maps with only minor inconsistencies were also found by earlier linkage mapping studies in *B. napus* (*12, 34*). The linkage map constructed here was on average 1.9× larger in cM compared to six published *B. napus* linkage maps generated using different approaches with relatively similar marker densities (*12, 14, 16, 34-36*). Although our mapping population is derived from two highly divergent parental lines and may enable us to sample more crossovers than other studies, some residual genotype errors or segregation distortion is expected to lead to some map inflation. In contrast to our study, which relied on an F_2_ population, five of the compared studies used recombinant inbred lines or doubled haploid populations, which will likely suffer less genotyping errors due to their inherent lack of heterozygous alleles. Using a genotyping array, which is less error-prone than ddRAD genotyping, in a *B. napus* F_2_ population also led to a smaller genetic map of roughly half the size (*16*). This suggests that ddRAD genotyping errors, and not the population type, are the main reason for the genetic map inflation. Nevertheless, the high-density and accuracy of the linkage map presented here suggest that it is useful for localizing QTL.

### Flowering time QTL on chromosome C2

Flowering time QTL in *B. napus* are mostly found in regions syntenic with Arabidopsis chromosome 5 (Parkin et al., 2005) on *B. napus* chromosomes A2, A3, A9, A10, C2 and C3. Here, we identified a single significant locus for flowering time and budding time on C2. These two phenotypes were significantly correlated, explaining the shared QTL that explained ∼9% of variance. This modest amount of variance explained is typical for traits that are controlled by many loci spread over multiple chromosomes, with each making minor contributions to the phenotypic effect. Additionally, the size of the QTL region (LOD confidence interval) differed for flowering time and budding time. The size of the QTL region is important, as it reflects the level of mapping resolution that was achieved. In F_2_ mapping populations, the confidence interval of a QTL can be large (>1Mb), and these populations often represent the starting point for fine-mapping of candidate genes. In this study we used the recommended LOD threshold of 1.5 units for 95% coverage of the confidence interval (*37*). However, the width of the confidence interval depends on how steep the QTL peak is, which can depend on a range of factors including marker density (*38, 39*).

The QTL overlaps with a locus identified in an earlier study, which found that the 60K array SNP Bn-scaff_18507_1-p889927 was associated with a QTL on C2 at position 33,936,984 on v81 (position on Darmor v41: 26,548,393) explaining 6.36% of flowering time variation (*26*). However, the QTL LOD peaks identified here are distant from this location. Among the known flowering time genes on C2, *FT* (*40*), *FLC* (*41*) and *FY* (*42*) may have particularly substantial effects. *FT* has been implicated in *B. napus* flowering time divergence (*28, 43*) and *FLC* has been found to explain ∼23% of flowering time variation in *B. napus* (*44*). *FY* is a suppressor of the transcription factor *FLC* (*42, 45*), and lies in close physical proximity to the LOD peaks identified in this study. However, the *FT* homolog on C2 is expressed at low levels in all ecotypes tested (*46*), which suggests it may not be the candidate gene. Further, none of the flowering time gene homologs occur within the budding time QTL region. Based on the QTL results of this study, we therefore cannot conclusively determine whether the C2 homologs and/or regulatory elements of *FY, FLC, FT* or another gene are underlying the detected QTL. The F_2_ experimental design is only the start of the discovery process because lack of recombination between closely linked regions can hinder high-resolution mapping. Our results show one significant QTL and additional suggestive regions of interest. It is likely that several of these loci would need to be transferred into the desired genetic background to exploit heterosis between spring and winter varieties.

### Genotype errors and correction

We detected high pairwise genotype discordance within the duplicate parental samples. Because genotype errors in either of the compared duplicate samples can lead to discordance, the genotype error rate can be roughly estimated as half of the discordance (∼6%). In line with GBS results in a rice F_2_ population (*47*), most errors can be attributed to undercalling of heterozygous alleles (97.51%). The parental lines are homozygous, though residual heterozygosity and mismapping can lead to heterozygous allele calls. The true error rate in the progeny is therefore likely higher than in the parents, because F_2_ populations contain an expected 50% heterozygous alleles.

Despite the range of optimal depths reported in the literature, it is clear that calling heterozygous SNPs accurately requires depths substantially higher than those required for calling homozygous SNPs (*48, 49*). The moderate sequencing depths used in this study (9.41×) may thus lead to inflation of sequencing noise and insufficient allele sampling, which can result in undercalling of heterozygous alleles (*50*). The percentage of errors attributed to undercalling of heterozygotes may even be an underestimate, as errors between apparently homozygous alleles may be caused by conflicting erroneous genotype calls of a heterozygous allele.

The genotype error rates found here are higher than error rates reported in the literature, even for heterozygous populations. For example, Malmberg et al. (2018a) analysed a heterozygous *B. napus* panel with different skim sequencing coverages and filtered genotype calls using a minimum read depth of 5. The authors found error rates of 2.1% error at 2 × sequencing coverage and 4.2% error at 1 × sequencing coverage (*50*). Similarly, an error rate of 3% was found using GBS in a bovine population with a minimum read depth of 5 (*51*). In a ddRAD genotyping study using a mapping population of cichlid fishes, an investigation of genotype errors found error rates of 4.41% at 8 × coverage (*52*). This suggests that GBS can lead to higher genotype error rates than expected in samples such as F_2_ mapping populations.

A substantial effect of genotype errors on linkage mapping was found. Cumulative inflation of linkage map length is often caused by genotype errors that introduce spurious double recombination events into the map (*53*). In addition, high levels of missing data and markers with segregation distortion may also affect the mapping distance as these alter the calculated recombination rate (*54*). It has been estimated that every 1% error rate in a marker adds approximately two cM to the linkage map (*55*). Linkage map inflation has been previously reported for GBS data in wheat (*53, 54*) and rice (*47*). In one of the studies on wheat, errors inflated the linkage by up to 11 times (*53*). In a linkage and QTL mapping study of *B. rapa* based on SNPs derived from GBS, high error rates (19.6%) were found and the resulting A genome linkage map was inflated, spanning 4,802.52 cM (*56*).

Linkage and QTL mapping in major crops are commonly carried out using highly accurate commercial genotyping arrays such as the Illumina Infinium Brassica 60K array (*57*). Genotyping arrays may introduce sampling bias because they only genotype previously known SNPs. A particular advantage of GBS over genotyping arrays is that regions missing from the reference genome can be genotyped and used for linkage mapping. However, genotyping arrays have the important advantages of more accurate calling of heterozygous genotypes and low missing data. Although genotype arrays also produce errors that frequently involve heterozygous sites, the error rates are likely lower at 1-2% (*58, 59*). Our results suggest that in heterozygous populations, genotyping arrays will generate markers with substantially higher accuracy than GBS. Here, to increase genotype accuracy for GBS data, genotype correction was applied. We find that genotype correction substantially decreased genetic map inflation, underlining the value of a correction step in heterozygous populations analysed using GBS at low to moderate sequencing coverage.

## Conclusion

We report a QTL on chromosome C2 for flowering time and budding time in a *B. napus* winter type *x* spring type cross. This QTL and the additional suggestive loci can be fine-mapped and backcrossed into the parental varieties to facilitate flowering time control in hybrid spring type *x* winter type varieties. An optimised combination of enzymes was also identified using in silico analysis, and the resulting number of empirical ddRAD loci and SNPs demonstrate the effectiveness of the enzyme combination HinfI and HpyCH4IV. In addition, we show that ddRAD generates high levels of genotype errors that can impact linkage map construction. By filtering SNPs and applying genotype imputation and correction an accurate map could be constructed, allowing effective QTL analysis of flowering time. Further investigation of the loci controlling flowering time and maturation will allow *B. napus* breeders to better exploit variation in winter and spring types.

## Materials and methods

### Plant material and phenotyping

The mapping population resulted from a cross between an early-flowering spring line (BnSOSR) and a late-flowering winter line (BnWOSR) carried out by BASF. The F_2_ population consisting of 200 individuals as well as 4 BnSOSR and 3 BnWOSR parental replicates was sown directly into 10×10×15 cm pots in a phytotron at the University of Western Australia, Perth. Temperature was maintained between 18-22 °C. To ensure flowering occurred, vernalization was initiated 63 days after sowing and plants were moved to a controlled environment chamber with a constant temperature of 4 °C and an 8-hour light period. Plants were watered twice a week by hand. After approximately 6 weeks, and a total of 108 days after sowing, the plants were returned to the phytotron. This 6 week vernalization period is sufficiently long to ensure that variation in flowering time in the F_2_ population is not driven by variation in vernalization response (*60, 61*). Each pot was provided with a dripper and connected to the irrigation system. Plants were watered twice a day for 1 minute. A total of 125 ml of fertilizer with micro minerals was provided by hand every two weeks. The time of first floral buds appearing and the date of first flower opening were recorded. All plants were grown until seed set. To investigate whether phenotype data was normally distributed, Shapiro-Wilke’s tests were carried and histograms plotted using ggpubr 0.2.1 (*62*).

### Restriction enzyme selection and digestion

DNA fragmentation was carried out by simultaneous digestion using two restriction enzymes. Suitable restriction enzyme pairs, that created sticky or overhanging ends, were selected based on their reaction buffer and incubation temperature compatibility to allow simultaneous digestion. The software IgCoverage 1.0 (*7*) was used to carry out in silico digestion of the *B. napus* genome (Darmor-bzh v8.1; *63*) using these enzyme pairs. The number of fragments within the 100-600 bp size range with different ends (LengthDeFrag100-600), as well as the expected percentage of genome coverage breadth generated by the selected 26 restriction enzyme pairs, were then compared. Out of the 26 suitable enzyme pairs selected, 18 pairs showed coverage breadth > 20% (Table S1). The enzyme pair HinfI and HpyCH4IV (New England Biolabs, Ipswich, USA) was selected based on the number of fragments, genome coverage, availability and cost per sample. This pair was predicted to generate 840,663 fragments with different ends within the 100-600 bp range, which covered 24.6% of the genome. The suitability of the selected restriction enzyme pair was confirmed by digesting 400 ng of genomic DNA using 5 Units of each restriction enzyme and NEB CutSmart™ buffer (10×) (New England Biolabs (NEB), Ipswich, USA). The reaction was incubated for four hours at 37°C and the results were visualised using the LabChip GX Touch 24 (PerkinElmer, Waltham, USA).

### Adapter design

Adapters for the ddRAD protocol were designed based on the adapters and indexed primers used by Peterson et al. (2012). Barcoded adapters were modified to create a complementary overhang for the HpyCH4IV restriction enzyme, while the common adapter was altered to create a complementary overhang for the frequent-cutter HinfI. The adapters were assembled by annealing 10 µM forward and reverse strand oligos as described in Peterson et al. (2012). The adapter concentrations to be used in the ligation step for the barcoded and common adapters were determined using the molarity calculator described by Peterson et al. (2012). The average distance between the restriction sites required for the calculation was calculated using the estimated in silico digestion results obtained using the IgCoverage package.

### ddRAD library preparation

Genomic DNA was extracted from leaf material using the DNeasy Plant Mini Kit (QIAGEN, Hilden, Germany) according to the manufacturer’s protocol. DNA concentrations were quantified using the broad range Qubit 3.0 Fluorometric assay (Invitrogen, Carlsbad, USA), while DNA quality was assessed with the LabChip GX Touch 24 (PerkinElmer, Waltham, USA). Modified versions of the Peterson et al. (2012) and Clark et al. (2014) protocols were used to construct the ddRAD libraries. The extracted gDNA was normalised at 50 ng/μL and 200 ng of DNA of each sample was digested in a 20 μL reaction volume containing restriction enzyme/s and recommended buffer. Digestion for the preparation of the ddRAD libraries was carried out using HpyCH4IV (5 U) and Hinfl (5 U) in NEB CutSmart™ buffer. The reaction was incubated at 37°C for 4 hours.

The digested DNA was ligated respectively to the unique barcoded and common adapters using T4 DNA ligase (Thermo Invitrogen, Carlsbad, USA). An 18 μL master mix containing ligation buffer, 200 U of T4 ligase and the common adapter was prepared and added directly to the 20 μL digest reaction, after which the individual barcoded adapters were added. The reaction was incubated at 22°C for two hours, followed by 65°C for 20 minutes, then cooled to 4°C at a ramp rate of 2°C per 90 seconds. To accommodate variation in DNA concentration and quality the samples were not pooled after ligation but individually purified and double size selected to enable enrichment of fragments between 250 bp and 800 bp. The total volume of the samples was adjusted to 100 μL by adding 60 μL of nuclease free water. Double size selection was carried out by adding 50 μL of a 1:4 (0.5X) mixture of AMPure XP Beads (Beckman Coulter, Brea, USA) to PEG buffer (20% PEG w/v, 2.5M NaCl) to remove fragments >800 bp. The supernatant was transferred to 20 μL of a 1:1 (0.7X) Ampure XP bead to PEG buffer mixture to collect fragments >250 bp. The beads were washed using 80% ethanol and the fragments eluted in 30 μL nuclease free water.

A 10 μL aliquot of the size selected DNA was used for PCR amplification. A 40 μL master mix of Phusion Hot-Start High-Fidelity Master Mix Polymerase (Thermo Fisher Scientific, Walthan, USA) and the Forward (0.5 μM) and Reverse primers (0.5 μM) was prepared. Samples were amplified at 98°C for 2 minutes, followed by 15 cycles of 98°C for 15 seconds, 62°C for 30 seconds, 72°C for 30 seconds, and a final extension for 5 minutes at 72°C. Amplified libraries were cleaned using 1.5X Ampure XP Beads to sample volume to remove primer dimers. The resulting library DNA concentrations were determined using the High Sensitivity (HS) Qubit 3.0 Fluoro metric assay. Library quality and fragment size distribution were visualised using the LabChip GX Touch 24. Equimolar amounts of the prepared libraries were pooled and loaded on a 1.5% agarose gel to enrich and select fragments between 300-700 bp. The DNA was recovered using the QIAquick Gel Extraction Kit (QIAGEN, Hilden, Germany). The final library concentration, quality and size distribution were assessed again and adjusted to 20 nM DNA using 10 nM Tris Buffer (pH 8.5, 0.1% Tween 20, 10 nM). The final libraries were sent to the KCCG Core facility at the Garvan Institute for Medical Research for paired end sequencing on the HiSeq X Ten platform.

### Adapter trimming and quality trimming

The Illumina bcl2fastq 2.20.0.422 pipeline (*64*) was used to convert base call files to FASTQ format. Paired-end sequencing reads were demultiplexed using sabre 1.0 (*65*) with a single mismatch allowed. Raw FASTQ files were trimmed of adapter sequences and low quality bases with Trimmomatic 0.36 (*66*). For adapter trimming, a maximum mismatch score of 2 was used for the adapter sequence, together with a palindrome clip score threshold of 30 and a simple clip score threshold of 10. Low quality bases with a Phred+33 score below 3 were trimmed from the start and end of the read. Sliding window trimming was carried out using a 4-base wide window, trimming the bases when the average quality per base was below 15. The Illumina TruSeq3-PE adapter list provided with Trimmomatic was used for adapter trimming. All reads with fewer than 36 bases after trimming were discarded. Reads that were unpaired after trimming were also discarded. Following read pre-processing, the untrimmed and trimmed reads were analysed using the diagnostic tool fastQC (*67*). The results of fastQC for each sample were then aggregated and summarised using multiQC (*68*). The multiQC report was used to verify that adapters had been removed and read quality was high.

### Aligning sequencing reads

Trimmed reads were mapped using BWA 0.7.17 with the BWA-MEM algorithm (*69*) to the *B. napus* Darmor-bzh v8.1 assembly (*63*) using default parameters. After alignment, SAM files were converted to BAM format using SAMtools 1.8 (*70*). Unmapped reads, supplementary alignments and reads with a mapping quality below 20 were discarded. This filter removes multi-mapping reads, which commonly occur in *B. napus* due to the homeologous regions of its polyploid genome. The mapping results were analysed using SAMtools stats and mosdepth 0.2.3 (*71*). The number of ddRAD loci were calculated from mosdepth per-base output using BEDTools 2.26.0 (*72*) to merge neighboring loci within 100bp.

### Calling single nucleotide variants

Variants were called using GATK 3.6 (*73*). First, BAM alignments were indexed using SAMtools, then HaplotypeCaller was used to call SNPs for each individual sample. Genotyping was carried out using GATK GenotypeGVCF using default setting with auto index creation and locking when reading rods disabled. Results per chromosome were merged using GATK CatVariants. Variants were filtered using VCFtools 0.1.15 (*74*). Indels and multiallelic SNPs were excluded *(--remove-indels --max-alleles 2 --min-alleles 2*). Before filtering SNPs individuals with >0.9 missing genotypes were removed. To reduce the rate of heterozygous alleles incorrectly called as homozygous alleles due to insufficient read depth, genotypes with a depth <5 *(--minDP 5*) were set to missing. SNPs were discarded if they displayed a minor allele frequency <0.05 (*--maf 0*.*05*) or when genotypes were not present in >80% of all individuals (*--max-missing 0*.*8*). Genotype discordance was calculated with snpEff 4.3t (*75*) using the duplicate samples for the parents (spring type BnSOSR with n=4, winter type BnWOSR with n=3) with pairwise comparisons of genotypes for SNPs passing the above filters. Heterozygosity per individual was calculated using VCFtools.

The parentage assignment and filtering of distorted SNPs was carried out using the custom script vcf2gt.py (*76*), which uses cyvcf2 0.8.0 (*77*) to parse VCF files. A chi-square test implemented in scipy 1.2.0 (*78*) was carried out to identify and discard SNPs with significant segregation distortion (p<0.01) based on the expected F_2_ segregation ratio of 1:2:1. Further filtering removed SNPs that were heterozygous in at least one of the parents or that were not polymorphic between the parents. The script also converted the SNPs in VCF format to a genotype matrix in AB format (A: homozygous allele from Parent 1, B: homozygous allele from Parent 2; AB: heterozygous allele; -: missing allele).

Genotypes were imputed and corrected using Genotype-Corrector 1.0 (*79*). This software uses the order of SNPs on the genome reference and a sliding-window approach to impute and correct genotypes based on neighboring genotypes in F_2_ populations. Before correction, up to eight consecutive homozygous SNPs within 150 bp genomic intervals in heterozygous regions were binned into a single SNP with Genotype-Corrector qc_hetero. This helps prevent miscorrection of heterozygous genotypes to homozygous genotypes when using the sliding window approach. With the 20% missing SNPs used here, the expected accuracy of Genotype-Corrector is >95%, based on empirical testing in crop mapping populations (*79*).

### Linkage mapping

Linkage mapping was carried out using the MSTMap algorithm (*80*) implemented in the R package ASMap (*81*). The sum of recombination events objective function was used to find the optimal sequence of loci, and the p-value threshold for clustering markers into linkage groups was set to 1e^-23^ based on evaluation of a range of values from 1e^-14^ to 1e^-29^. The kosambi distance function was used to estimate genetic distances between SNPs, and rare recombination events were treated as errors (*detectBadData=True*). Linkage groups were assigned chromosome names based on marker positions on the reference genome. Small linkage groups that did not represent an entire chromosome, were merged with chromosomal linkage groups and genetic distances recalculated if an unambiguous assignment was possible using physical marker positions. Linkage groups with <7 markers were discarded. The estimated pairwise recombination fractions between markers were calculated using the rqtl function plotRF (*37*). Recombination fractions were used to identify outlier markers that are not in LD with neighboring markers and to manually correct marker order using physical marker positions. A total of 95 outlier markers were removed from further analysis. To ascertain the quality of the genetic map, the correlation between marker order on the genetic map and the reference genome was calculated using a Spearman’s rank correlation test in R. Crossover frequency was estimated using the rqtl function locateXO and a custom python script crossover.py (*76*).

### QTL mapping

QTL mapping was conducted with rqtl scanone using a single QTL model and the non-parametric model for flowering and budding time because these traits did not follow a normal distribution. Genome-wide significance thresholds for logarithm of the odds scores (LOD) were estimated using a permutation test with 1,000 iterations (*82*). The 1.5-LOD drop interval of each QTL position was estimated using rqtl lodint. The percentage of variance explained for each QTL was calculated using rqtl fitqtl with Haley-Knott regression. The custom interactive R script used for QTL mapping in Rstudio 1.1.456 (*83*) was based on the rqtl manual (*37*).

### Flowering time genes

A total of 306 Arabidopsis flowering time (FT) genes from the FLOweRing Interactive Database (*84*) was downloaded from The Arabidopsis Information Resource (*85*). These genes include homologs of the known *B. napus* flowering time genes. BLAST+ 2.2.29 (*86, 87*) analysis of the FT genes against the reference genome was carried out to find gene homologs using a cut-off of 1e^−6^ (following *88*). Overlapping hits were merged using BEDtools. The gene names in the v81 annotation (*63*) were identified using BEDtools to obtain gene annotations overlapping the BLAST alignments.

## Acknowledgements

A.S. was supported by an IPRS awarded by the Australian government. This work was supported by the Australia Research Council (Projects LP160100030, LP140100537 and LP130100925) and BASF. Bayer transferred this work with the CropScience business to BASF in 2018. This work was supported by resources provided by the Pawsey Supercomputing Centre with funding from the Australian Government and the Government of Western Australia.

## Author contributions

DE, JB and SR conceived and supervised the project. ASE grew the plants and carried out phenotyping. ASE, AP and DP prepared the sequencing libraries. ASE wrote the phenotyping and library preparation sections of the Materials and Methods. AS carried out the analyses and drafted the manuscript. AS, DE, JB and SR wrote the final manuscript and all authors provided comments and edits.

## Data availability

All sequences have been deposited in SRA (PRJNA640838) with individual accessions listed in Table S8.

## Code availability

Scripts used to for this study are available at https://github.com/ascheben/bn_gbs/.

## Supplemental tables

Table S1. In silico double enzyme digest analysis. Frag: Total number of fragments; FragDe: total number of different end fragments; LengthDeFrag: total length in bases of different end fragments; DeFrag100-600: total number of different end fragments between 100 bases and 600 bases; LengthDeFrag100-600: total length in bases of different end fragments between 100 bases and 600 bases; % Coverage: percentage of the reference genome covered by fragments between 100 bases and 600 bases. https://github.com/ascheben/bn_gbs/blob/master/supplemental_tables/Table_S1.xlsx

Table S2. Linkage map constructed using ASMap with corrected and imputed ddRAD markers derived from an BnSOSR x BnWOSR F2 population. https://github.com/ascheben/bn_gbs/blob/master/supplemental_tables/Table_S2.xlsx

Table S3. Mean markers and linkage map sizes across maps generated in this study. Abbreviations: doubled haploid (DH), recombinant inbred lines (RIL), amplified fragment length polymorphisms (AFLP), Brassica 60K genotyping array (Brassica 60k). https://github.com/ascheben/bn_gbs/blob/master/supplemental_tables/Table_S3.xlsx

Table S4. Phenotypic variation in flowering time and time to bud in the F2 mapping population and the parental lines. Phenotypes of spring type BnSOSR and winter type BnWOSR parental lines are shown as means of all replicates of each type (n=4, n=3, respectively). https://github.com/ascheben/bn_gbs/blob/master/supplemental_tables/Table_S4.xlsx

Table S5. QTL identified using genome-wide single QTL scans for budding time (B) and flowering time (FT). For each QTL, the phenotypic variance explained (PVE) and additive effect (AE) are shown. https://github.com/ascheben/bn_gbs/blob/master/supplemental_tables/Table_S5.xlsx

Table S6. Candidate flowering time genes within the QTL interval detected on chromosome C2. https://github.com/ascheben/bn_gbs/blob/master/supplemental_tables/Table_S6.xlsx

Table S7. Phenotype data for flowering time and budding time in the mapping population and parents. https://github.com/ascheben/bn_gbs/blob/master/supplemental_tables/Table_S7.xlsx

Table S8. Sample list for the 192 progeny and 7 parental individuals used in this study. https://github.com/ascheben/bn_gbs/blob/master/supplemental_tables/Table_S8.xlsx

## Supplemental figures

**Figure S1.**
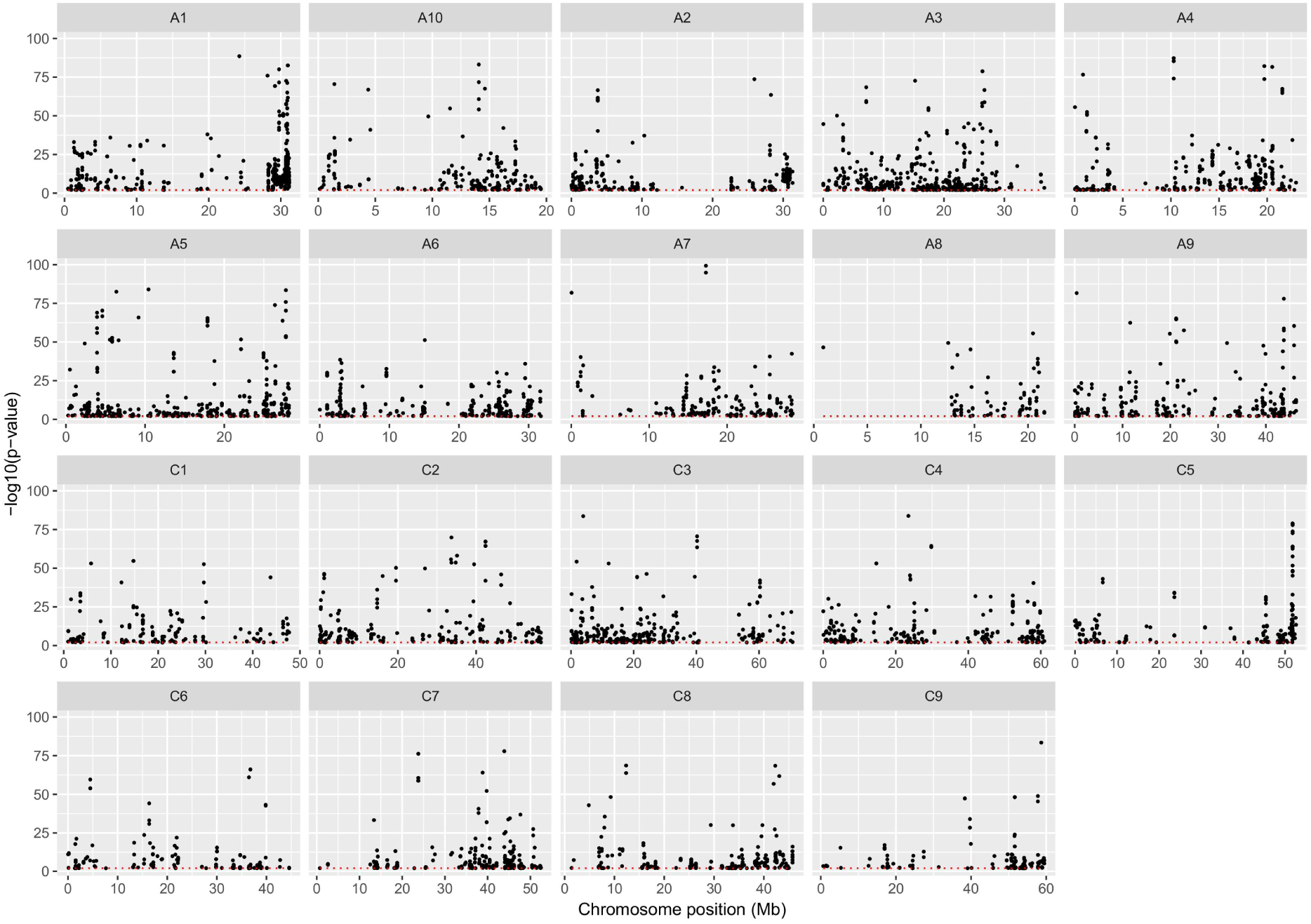
Segregation distorted loci across all chromosomes. The significance threshold (p>0.01) is shown as a red dotted line.

**Figure S2.**
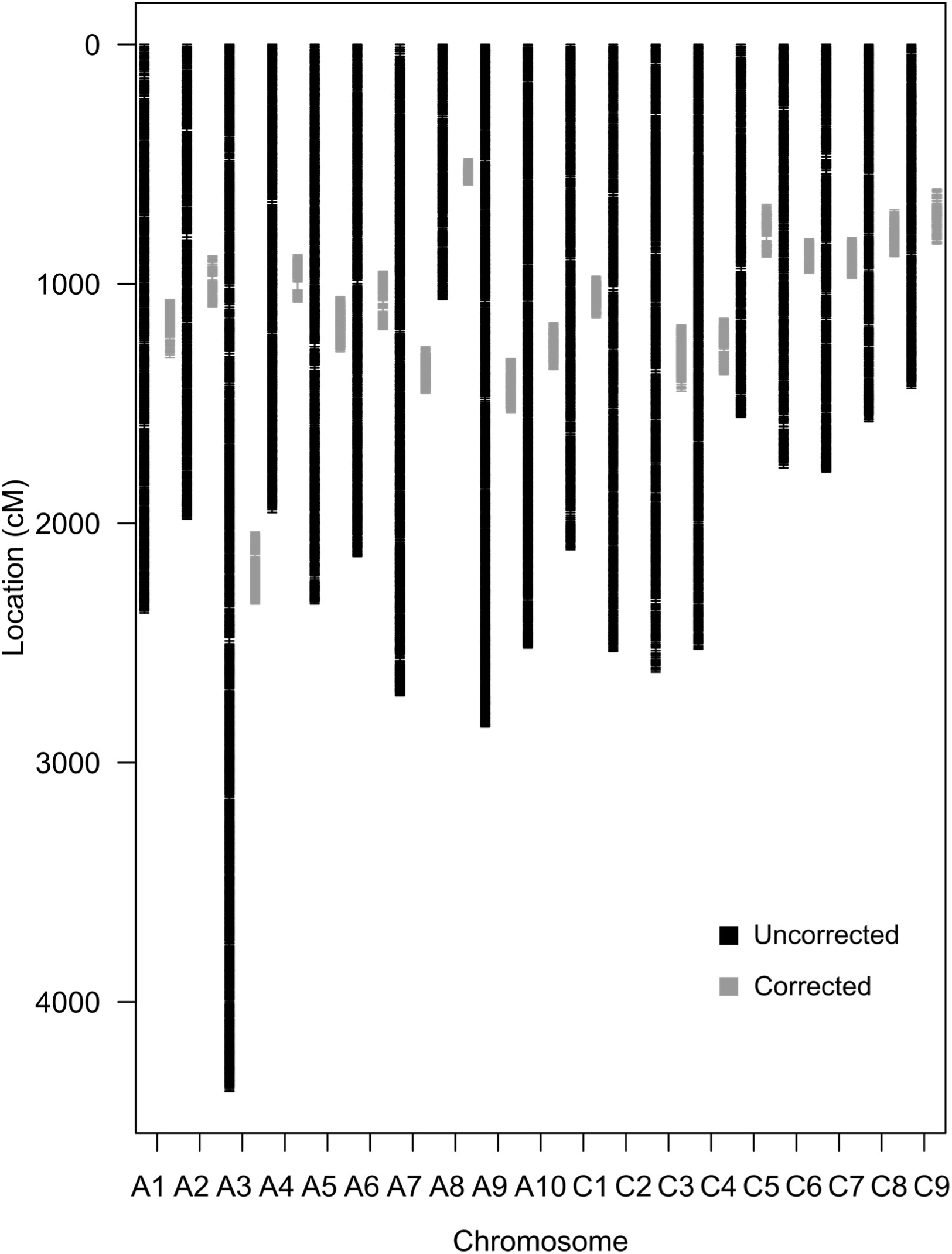
Comparison of linkage group sizes in an uncorrected genetic map and a corrected genetic map. Corrected linkage groups are aligned centrally to uncorrected groups.

**Figure S3.**
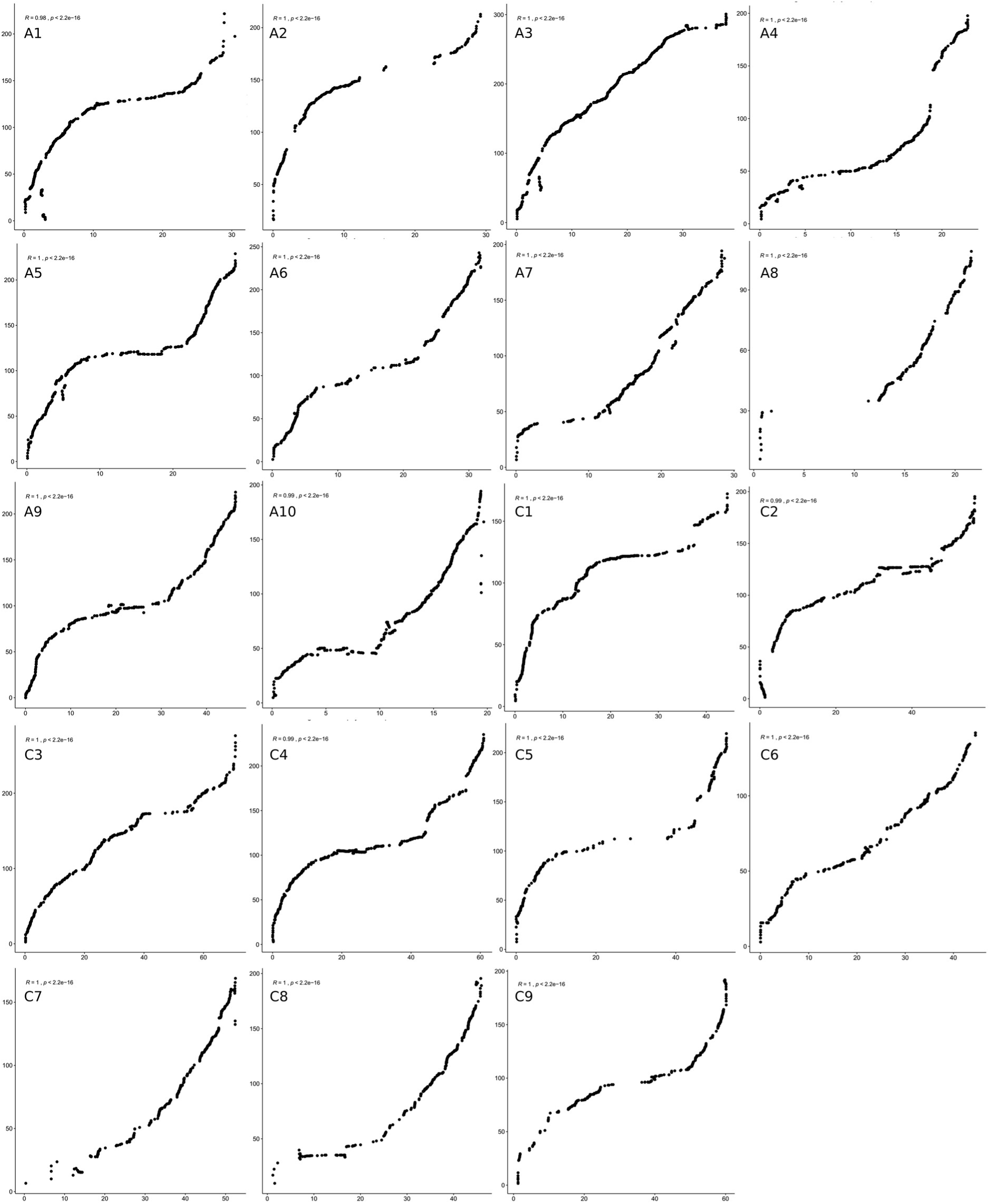
Physical (x-axis) and genetic (y-axis) marker positions on all chromosomes in Mb and cM respectively. Spearmans’s rank correlation test result shown in the top left corner of each plot.

**Figure S4.**
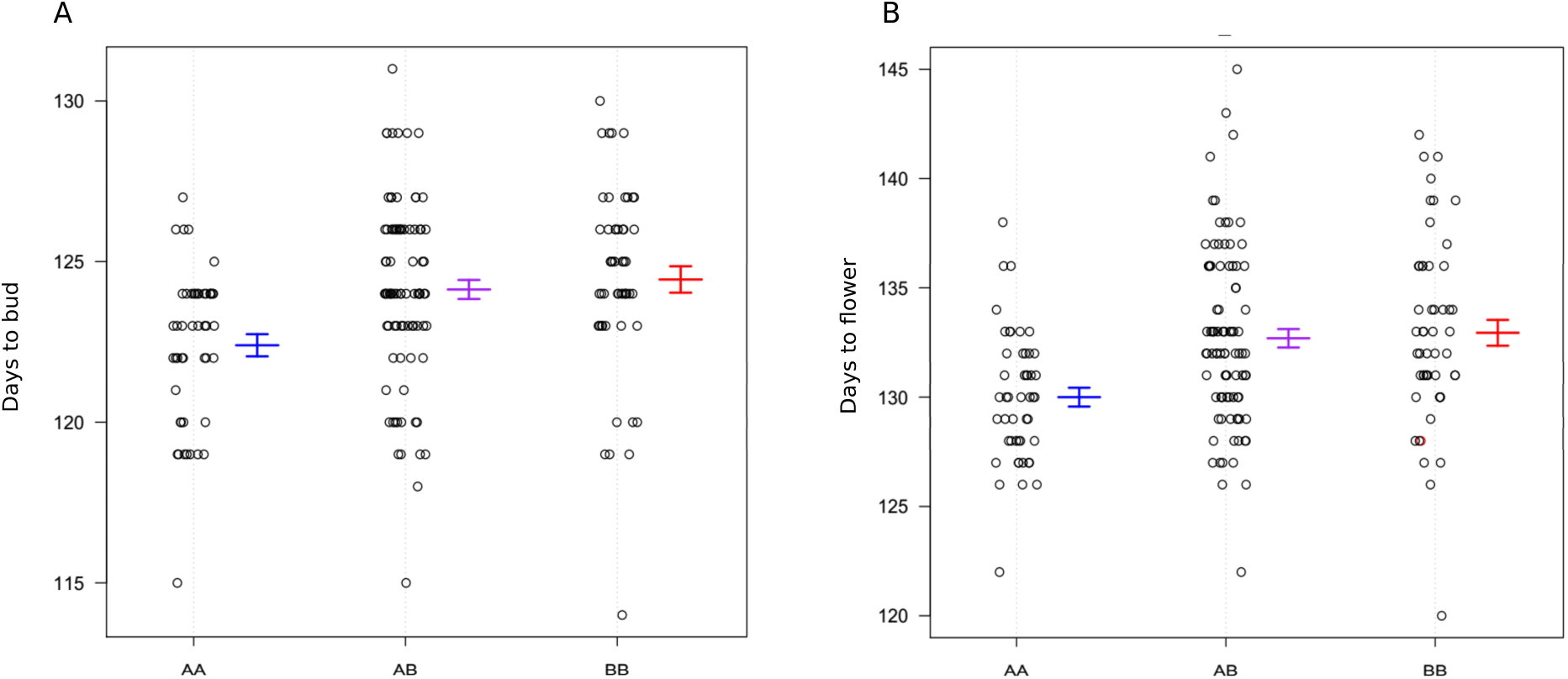
Effect plot for the budding time (A) and flowering time (B) QTL on C2 at positions 4,673,904 and 4,655,461 respectively. The ‘AA’ genotype is BnSOSR and the ‘BB’ genotype is BnWOSR.

